# Physics of low complexity polymers: patterning, polymer collapse and bulk microphases

**DOI:** 10.1101/2025.11.21.689704

**Authors:** Martin Girard

## Abstract

Polymer physics has arisen as a pillar of understanding behavior of intrinsically disordered proteins. Behavior of polymers can be understood in terms of generic behavior. Specifically, phase behavior can be understood in terms of solvent quality and chain length. The theory is well established for homopolymers, and specific heteropolymer structures, in particular random and co-block polymers. However, protein sequiences are neither of these. Nevertheless, there is a clear link between sequence and properties, which has given rise to the concept of molecular grammar.

In this article, we establish a framework to understand the behavior of Sturmian polymers, which provide a reasonable non-random polymer model for disordered proteins. Specifically, we show that molecular grammar arises as a non-linear critical susceptibility near the *θ* transition of single chains. We demonstrate that molecular grammar exists in polymer melts, where is determines a length scale under which chains are compressed, and above which chains are expanded.

## INTRODUCTION

The idea that protein condensates formed by liquidliquid phase separation is now prevalent in biology. Such condensates would provide means to spatially organize cells and provide distinct environments, e.g. for specific chemical reactions to occur [1]. A significant fraction of proteins thought to drive condensate formation possess intrinsically disordered regions (IDR), which are often prion-like low-complexity regions [2]. In turn, this has led to a large amount of research devoted to understanding the link between IDR sequence and phase separation, now termed molecular grammar [3].

From a physics perspective, behavior of a polymeric system is intrinsically tied to the behavior of a single chain [4, 5]. Specifically, phase separation into polymer-rich and -poor phases is only possible below the so-called *θ* temperature, which corresponds to the coil-to-globule transition of single chains. This naturally implies that quantities such as the radius of gyration (*R*_*g*_) of single chains is directly tied to the phase separation of multiple chains, which has indeed been confirmed for disordered proteins [6]. Unlike homopolymers, the sequence of monomers in a heteropolymer plays a role, and it has clearly been demonstrated that clustering aromatic or charged residues tends to collapse single polymer chains, and promotes phase separation [7]. From a theoretical perspective, approaches based on random phase approximation have been proposed to quantify this behavior [8, 9], yielding the so-called sequence decoration parameters.

Recently, we have developed a surrogate polymer model for low-complexity sequences, termed Sturmian polymers [10]. The procedure is based on combinatorics on words [11], which allows us to procedurally generate sequences with specific balance, which takes the role of block size. By investigating the behavior of single chains, we have observed that sequence specificity appears to be tied to the *θ*-transition. Specifically, for a given set of sequences with identical average (mean-field) hydropathy, dispersion in observed values of *R*_*g*_ is only observed near the polymer collapse transition. Within this region, the sequence hydropathy decoration was also found to be a poor predictor. This raises an interesting question: is molecular grammar a feature of the *θ* transition?, and if so, does it play any role in polymer melts?

Here, we provide strong evidence towards an a−irmative answer to the link between *θ* transition and molecular grammar. We first develop a non-linear susceptibility framework, in which molecular grammar can be reliably extracted as a second-order susceptibility. This provides molecular grammar with a rigorous quantiative thermodynamic framework. By performing finite-size scaling analysis, we show that non-linear susceptibility can be reasonably collapsed into a master curve with exponents compatible with the *θ* transition. Through scaling arguments and numerical experiments, we show that this susceptibility is always negative, resulting in low-complexity heteropolymer chains being more compact than their homopolymer counterparts. In polymer melts, the structure is affected, with small chains being compressed, and long chains being expanded with respect to a homopolymer melt, hinting at the presence of a bulk microphase transition.

### I. THEORY AND MODEL

#### Polymer model

In the context of this paper, we are interested in simple bead-spring models. These generally provide good approximation of interactions for intrinsically disordered proteins [12–14]. We denote the polymerization degree by *N*, and assume that all non-bonded interactions take place through an Ashbaugh-Hatch potential:

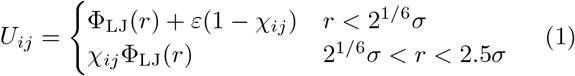

Where *ε* sets the energy scale and Φ_LJ_(*r*) is the usual Lennard-Jones potential:

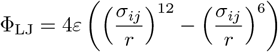

For heteropolymers, we use the combination rule 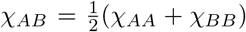 and 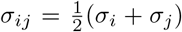. As previously discussed in [10], in the mean-field regime where all beads are well mixed and no correlations exist, the mixture should behave as-if all beads had mean-field Flory parameter *χ*_*e*_ = ⟨ *χ* ⟩ . Chain connectivity is enforced by harmonic bonds with spring constant *k* = 1000*ϵ/σ*^2^ and equilibrium distance *r*_0_. This distance is taken to be *r*_0_ = 1.1*σ*, which results in bond crossing, absence of entanglements, and fast relaxation dynamics. Absence of entanglements also results in avoidance of the glass transition in polymer melts. However, this does not fully avoid relaxation issues, such as Rouse time scaling 𝒪 (*N* ^2^), and generally slow dynamics at high densities. Consequently, the reader should be aware that statistical quality of simulation tends to decrease for long chains (see Fig S1 and associated discussion).

##### θ-point and the collapse transition

In the context of this article, we use the parameter 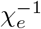 as a temperature. The reduced temperature in the context of polymer collapse therefore reads as *τ* = (*χ*_*θ*_*/χ*_*e*_ −1), where *χ*_*θ*_ is the interaction parameter at which the chains are in *θ* conditions. While this does not affect the transition itself (near *τ* ≈ 0), it is worthwhile to note that it is not strictly equivalent to a temperature. We determined *χ*_*θ*_ ≈ 0.416 based on finitesize scaling analysis of 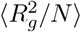 versus *τ N* ^1*/*2^ (see Fig. S2). This value is consistent with polymer collapse being observed near *N* = 1024 for *χ*_*e*_ = 0.45.

#### Reweighting and non-linear response

Consider a potential conformation 𝒞 and its associated statistical weight *w* = exp(− *βE*) of a homopolymer. We now introduce some disorder 𝒟, with associated magnitude *δ* in the homopolymer. The particular type of disorder is left here unspecified as the framework is generic, but we will later associate it with heteropolymer sequence. Our initial homopolymer conformation now has a statistical weight *w*_𝒞,*δ*_ = exp(− *βu* _𝒞,*δ*_), where *u* _𝒞,*δ*_ is the change in internal energy induced by the disorder. For clarity of the derivation, we assume that it is specified in thermal energy units, so that *β* = 1. We now expand the internal energy into a series:

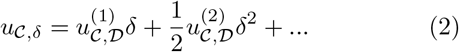

Where 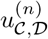 is the *n*-th partial derivative of the internal energy with respect to the disorder magnitude. We now turn to the standard reweighting expression for any observable *A*:

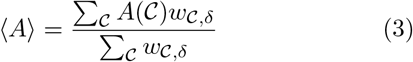

As with the internal energy, we now express the reweighted observable ⟨*A*⟩ as a series near *τ* = 0:

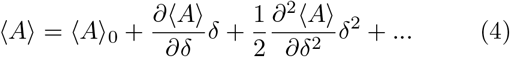

The partial derivatives of the above equation represent the non-linear susceptibilities of the chain with respect to disorder 𝒟. Specifically, we use the notation 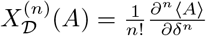 based on standard non-linear susceptibility notaion. *⟨* A *⟩* is expressed as:

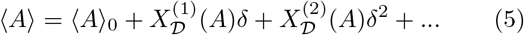

These susceptibilities take the form of covariances, as usually expected from statistical mechanics. They can be derived by substituting 2 into 3, and evaluating the partial derivative with respect to *τ* near *τ* = 0. The first (linear) susceptibility takes the usual form:

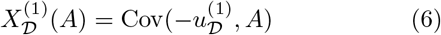

Whereas the second order susceptibility takes a slightly more complex form:

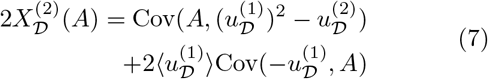

For the purposes of this article, the main observable of interest is the radius of gyration *R*_*g*_. For the sake of simplicity, any susceptibility with respect to unspecified observable explicitly refers to *R*_*g*_, such that 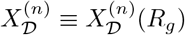.

#### Sequences and hydropathy decoration

Within this article, we consider disorder in terms of variation of the *χ* parameter along the chain length. Specifically, we consider binary sequences produced by Sturmian sequences and concatomers of Sturmian sequences (see [10]) of balance ℵ. We label this specific disorder with Δ, so that any term in *δ* in equations 2 - 7 is now replaced by Δ to indicate its specificity. The *χ* parameters are set to *χ*_*AA*_ = *χ*_*e*_ + Δ*/f*, and *χ*_*BB*_ = *χ*_*e*_ − Δ*/*(1 − *f* ), where *f* is the fraction of A beads in our Sturmian sequence. This particular transformation leaves the mean energy in the system unchanged according to the mean-field, i.e 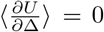. The partial derivative *u*_𝒞,Δ_ takes a simple form over pairs of parti-cles:

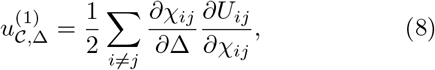

and any higher order derivative is zero since the potential (eq 1) is linear in *χ*_*ij*_.

During sequence generation of Sturmian polymers, we require that for given set of slopes, the balance for *N* = 64 and *N* = 1024 to be equal. This ensures that the observed balance is consistent for any *N* . Concatomers are based on Sturmian polymers of length *N* = 64 where we only require the sequence to be idempotent.

## II. SINGLE CHAINS

We first investigate single chains at infinite dilution. For these chains, we can make scaling arguments to predict general behavior. Specifically, consider for a moment that our homopolymer conformation is approximately a mixture of beads with density 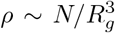, with no correlation related to the chemical distance along the backbone. Further consider that this results in contact between beads being random, independent and identically distributed. This allows us to relate partial derivative 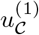 as being a random variable produced by the sum of partial derivatives of contact energies.

Given that we are assuming random contacts, the probablity of a contact is proportional to the product of fractions, i.e. the probability of an AB contact is *p*_*AB*_ = *f* (1 − *f* ). This allows us to write the variation of energy of a single contact to be:

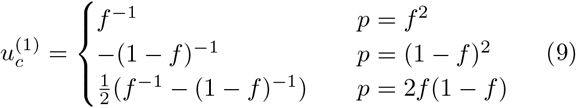

with 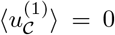, independently of *R*_*g*_, and therefore we have 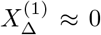. Practically, one would expect finite values from polymer ends, and local correlations in the sequence breaking the random contact approximation. Indeed, we observe small but non-zero values in Fig. 1A. These results suggest that the second order susceptibility can be well approximated by:

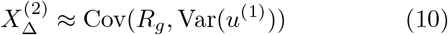

**Figure 1.**
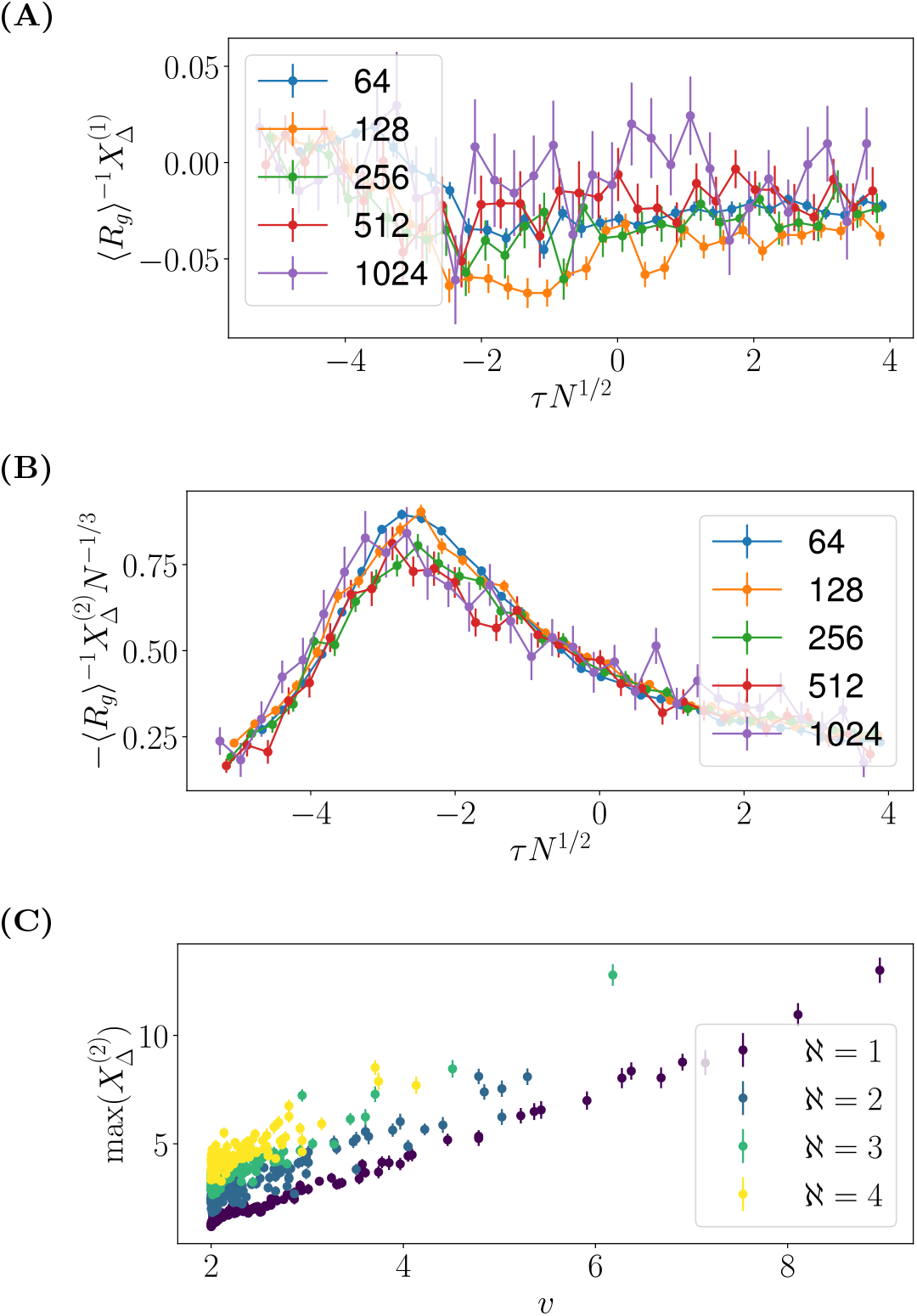
Finite-size scaling analysis of normalized second order susceptibility of polymer chains near the collapse transition, shown here for a particular sequence with ℵ = 1. Other examples are shown in the SI.

This approximation allows us to easily make predictions; the variance of *u*_Δ_ will increase with the number of contacts, which itself decreases with *R*_*g*_. It therefore follows that 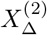 should be strictly negative for any sequence, bringing us towards a universal rule: at constant mean-field potential, any heterogeneity in the sequence decreases the radius of gyration. This prediction can also be validated by simulations, as shown on Fig 1B. Furthermore, based on eq 9, 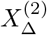 should be directly proportional to 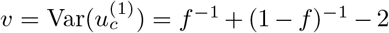, which is also numerically verified in Fig 1C.

### The θ transition

We now turn to the question of sequence specificity; namely whether it is directly tied to the polymer collapse transition. This stems from an observation made in [10], where dispersion of data across sequences is the largest at the chain collapse. In order to answer this question, we turn to finite-size scaling. Specifically, we investigate the relative second-order susceptibility,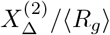, as a function of the rescaled temperature *τ N* ^−1*/*2^. When the susceptibility is rescaled by *N* ^−1*/*3^, we obtain approximate collapse of the curves as shown in Fig. 1B, with a peak near *τ N* ^−1*/*2^ ≈ − 2.8. This cements sequence-specificity to be intrinsically tied to the polymer collapse transition. It should also be pointed out that our polymer exhibits phase separation at *τ N* ^1*/*2^ ≈ − 1, indicating that the peak of susceptibility is not directly linked to phase separation. We provide a justification for the large exponent in the next section, although not for its precise value. The reader should keep in mind that goodness of the collapse is not perfect, and that the specific value of the rescaling exponent (1/3) should be viewed with skepticism.

### A note on mean-field transformations

It should be noted that 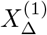 only vanishes for null mean-field transformations; If we consider instead a change *χ*_*AA*_ = *χ*_0_ + *a*Δ, *χ*_*BB*_ = *χ*_0_ + *b*Δ, then eq 9 would yield ⟨*u*^(1)^ ⟩ ∼ *af* + *b*(1 − *f* ), which represents precisely the change in *χ*_*e*_ = ⟨*χ* ⟩. In the case of disordered proteins, it is always possible to express their properties in a sequence-independent *χ*_*e*_ plus a sequence dependent *X*^(2)^.

#### Range of validity and change of *χ*_*θ*_

It is worthwhile to investigate the range of validity of the series in eq. 5. In order to do so, we simulated chains with *f* ≈ 0.15, and ℵ ∈ [1, 3], *N* ∈ [64, 256]. These values are chosen to be somewhat representative of prion-like disordered proteins. The data in Fig. 2A clearly shows that the expansion is valid until Δ ≈ 0.2. This may appear as a small value at first glance; however we have approximately *χ*_*A*_ ≈ 1.75 and *χ*_*B*_ ≈ 0.18, which is an extremely strong contrast between stickers and spacers. When compared to *χ*_*θ*_, the stickyness of the stickers is largely above the typical values found in force-fields for disordered proteins such as calvados [12] or MPIPI [13].

**Figure 2.**
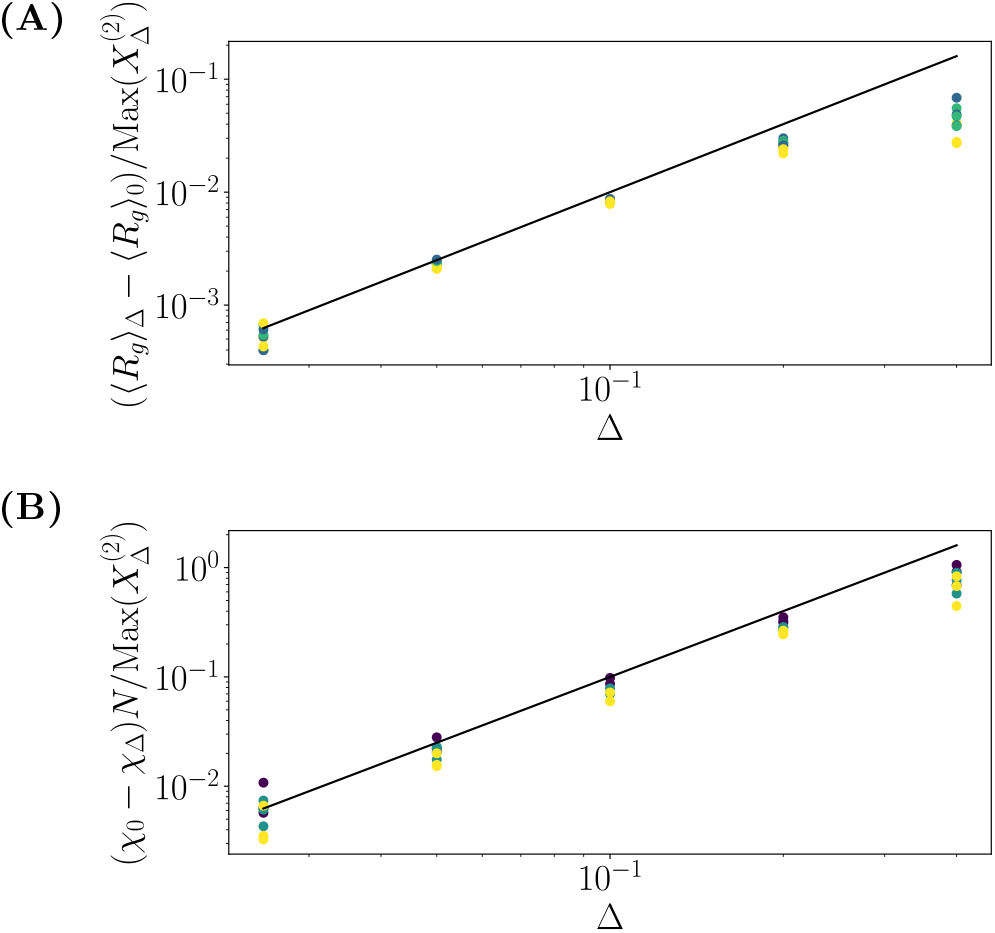
Chains with sticker-spacer architectures. Here, *N* ∈ [64, 256], ℵ ∈ [1, 3] and *f* ≈ 0.15. a) Changes in the radius of gyration versus Δ, normalized by the non-linear susceptibility. Black line indicates series expansion prediction, i.e. Δ^2^. Deviations are observed for Δ ≳ 0.2. b) Changes in *χ*_*θ*_ induced by disorder, measured by 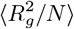 (approximate). Uncertainties are not displayed.

Introduction of disorder into the sequence can be seen as a shift of *χ*_*θ*_. In other words, the transition itself is left unchanged, but pushed to higher temperatures. Estimating *χ*_*θ*_ for chains with Δ *>* 0 with *N* ∈ [64, 256] poses significant challenges. We characterize *χ*_*θ*_ by measuring the shift in *χ* parameter corresponding to a fixed value of 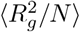, here chosen to be 0.3. Because of finite-size effects for small *N*, this measure is approximative. However, *χ*_*θ*_ clearly shifts towards higher temperatures, with the same underlying scaling (Δ^2^), and a similar range of validity. The reader should note that there is no proper theory to justify this observation; the *θ* temperature is not an observable of the system, and cannot be cranked through the reweighting framework of eq. 3. In addition, the scaling *N* ^1^ observed here is different than would be predicted based on Fig. 1, which would be of *N* ^0.83^.

## III. POLYMER MELTS

We now turn to polymer melts at fixed density. We specify melts by their bead volume fraction *ϕ*. Given our simple model, the rest of the volume would be filled by solvent. However, the reader should be reminded that a closed packed structure has volume fraction *ϕ* ≈ 0.74, so that this model can never yield a polymer-only phase. Here, we investigate volume fractions *ϕ* ∈ [10^−2^, 0.5], with particular attention in the regime *ϕ* ∈ [0.1, 0.5]. Polymer lengths are *N* ∈ [64, 512].

The susceptibility of chains in the melt is shown in Fig. 3A for *N* = 64. The high compression at low values of *ϕ* is consistent with single-chain observations. As density is increased, susceptibility goes through a maximum, located near *ϕ* ≈ 0.3. Interestingly, the dependence of *X*^(2)^ on temperature appears to be minimal at high volume fraction, independently of *N* (see SI for *N* = 128). At low density, the susceptibility is inconsistent with single chain data below *τ N* ^1*/*2^ ≈ −1, which is due to phase separation.

**Figure 3.**
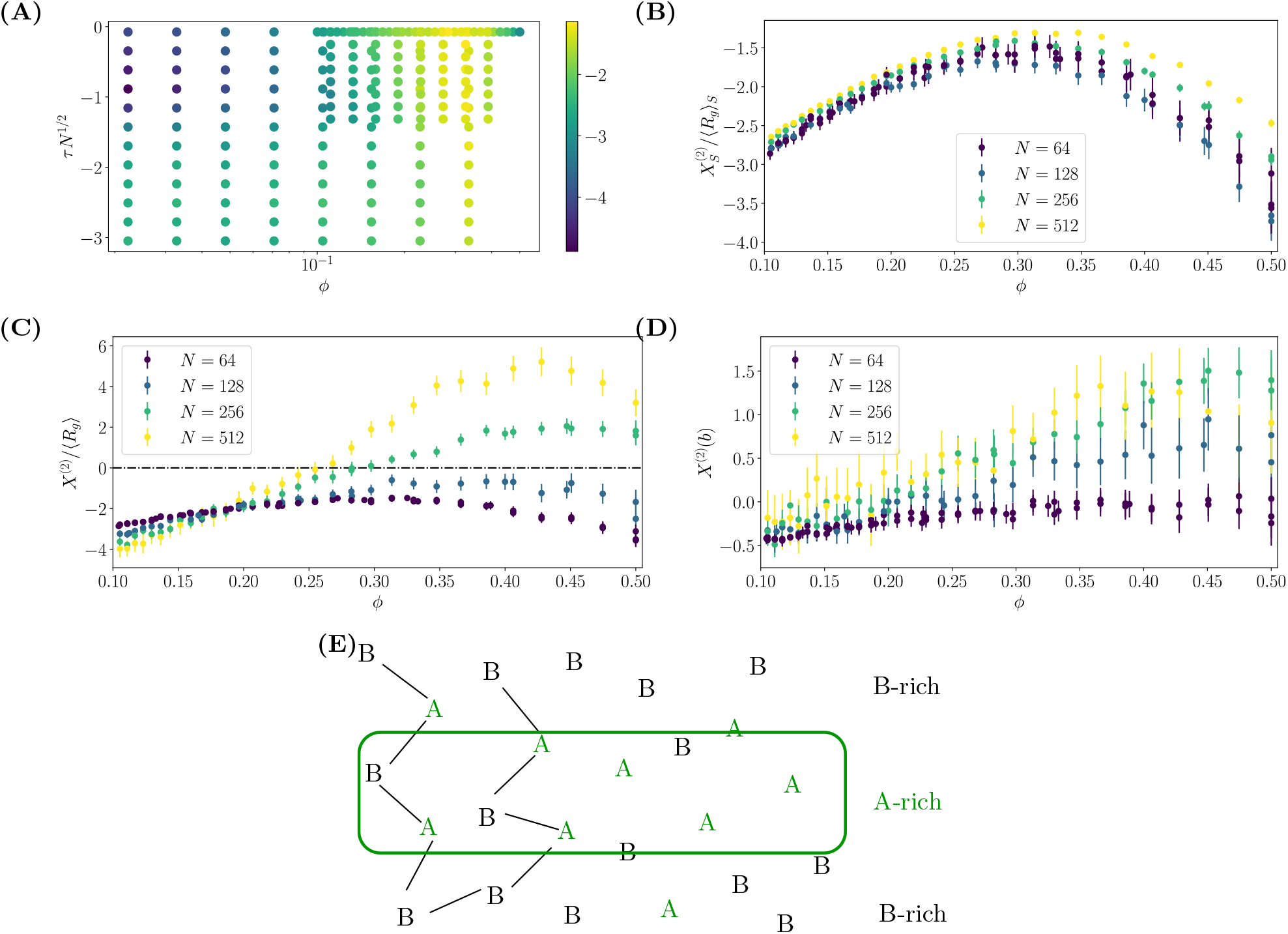
Second-order susceptibility of a polymer melt in constant mean-field. Results are presented for a sequence with *f* ≈ 0.23 is used for calculations. A) Second-order suscepbility for *N* = 64 versus volume fraction *ϕ* and reduced temperature *τ N* ^1*/*2^. Values at low density are inconsistent with single-chain data due to phase separation. For high density, the resulting susceptibility is only weakly dependent on temperature. B) Second-order susceptibility for segments of size *Ns* = 64 taken from melts of polymer with length *N* . The curves are in qualitative agreement, with differences likely arising from chain end effects, and lack of high quality data in high density melts (see SI). All segments are compressed by disorder. C) Second-order susceptibility of whole chains in the same melt. Curves are in qualitative disagreement, with longer chains being stretched by disorder. D) Second-order susceptibility of the polymer asymetry of whole chains in the melt. Short chains are more spherical than homopolymers, whereas longer chains are more elongated. Error bars indicate standard error across chains. There are two sets of data for *N* = 64, with 256 and 512 chains. E) Hypothesized worm-like chain microphase arising at high Δ. The core of the worm-like chain is depicted by a green rectangle, and it is enriched in A monomers (stickers). Locally, the polymer backbone (black lines) needs not follow to the worm-like chain direction. This leads to distinct signs of the susceptibility being observed at different length scales.

Given this independence with respect to temperature, we now examine properties of the chains at *χ* = 0.42. We first examine the properties of short (*N*_*s*_ = 64) segments. The susceptibility is shown in Fig 3B, and the results for different *N* are in general agreement, with small differences likely arising from the small differences in reduced temperature *τ N* ^1*/*2^.

Results for whole chains are shown in Fig 3C. It is clear that the curves for different polymer lengths are not in agreement, particularly with respect to the sign of the susceptibility. For *N* ≥ 256 and large *ϕ*, the susceptibility is positive indicating that although short segments are compressed, the whole chain adopts expanded conformations. Since single chains are always compressed by disorder, this effect clearly arises from collective behavior of the melt.

We hypothesize that this behavior is related to bulk microphases. The stickers want to be packed as tightly as possible to maximize contacts, expelling spacers from their local environment. The latter are however chemically bonded. One potential geometry to alleviate this frustration is to form a worm-like chain, with stickers at the center and spacers on the outside (sketched on Fig 3E). Although the polymer is flexible, the final geometry would still exhibit stiffness on longer length scales. Inter-chain repulsion in such a configuration is enhanced through local spacer concentration, resulting in lower overlap between chains than in a homopolymer. To futher support this argument, we turn to polymer asymmetry, defined by 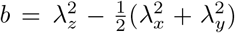, where *λ*_*i*_ are the eigenvalues of the normalized gyration tensor (*λ*_*z*_ *> λ*_*y*_ *> λ*_*x*_), 0 *< b <* 1. Applying the framework for non-linear susceptibility, we obtain *X*^(2)^(*b*) as shown in Fig 3D. For short polymers, or at low volume fractions, the chain is more spherical than a homopolymer (*X*^(2)^(*b*) *<* 0). Conversely for a long polymers at high volume fraction, the chain is found to be more extended than homopolymers. This picture supports our wormlike chain bulk microphase picture, although it is not conclusive.

Molecular grammar therefore exists in the polymer melt, but takes a different role than in single chains. Specifically, it determines the length scale at which the chain is “long”. For segments of length *N*_*s*_ = 64, the susceptibility is similar to single chain results (see Fig 4A). Namely extreme values of the sticker fraction, and blockiness determine the susceptibility. However, the profile for whole chains is flattened (see Fig 4B) at a positive value, and only extreme values of the sticker fraction shows any hint of sequence susceptibility. For smaller sticker fractions, e.g. *f* ≈ 0.08, longer chains are required to observe extension from disorder (see Fig. S4).

**Figure 4.**
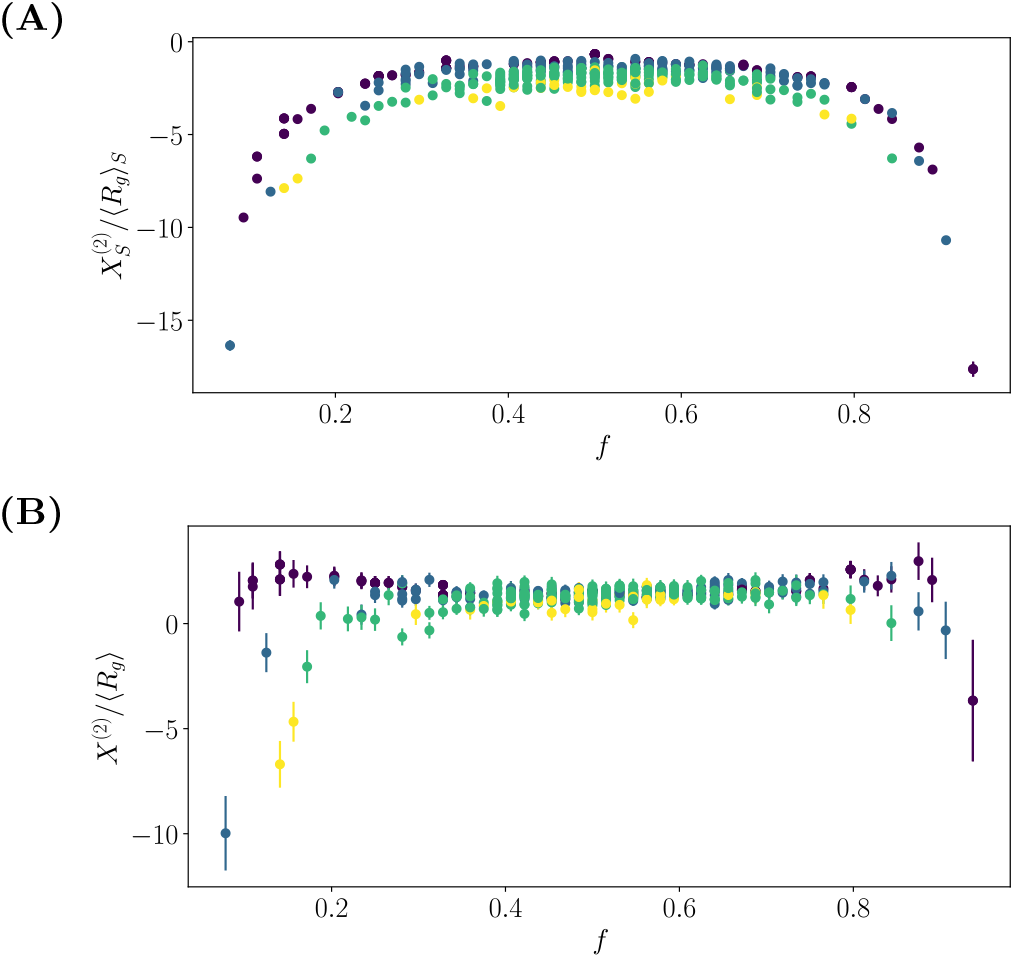
Representation of the molecular grammar for melts with *N* = 256 for A) Segments of length 64 and B) Full chains. Color indicates sequence balance. Error bars represent standard error across the different chains in the polymer melt

## IV. DISCUSSION

We have shown that sequence patterning is not a simple change of radius of gyration, rather it is a change in transition temperature, and is an intrinsically non-linear phenomenon. Specifically, it appears to be tied to the *θ* point in the same fashion any susceptibility would. This implies tools available near critical points are perfectly suited to further understand patterning in polymers. The large exponent observed for susceptibility ≈ 0.83, should however be interpreted as crossover phenomenon. Indeed, disorder affects the *θ* temperature as shown in Fig 2, which results in the chains being pushed from an ideal to a poor solvent regime. The exponent associated with this shift *N* ^1^ is itself larger than the exponent associated with *X*^(2)^. However, we do not have any theoretical justification for the observed values.

In polymer melts, polymer conformations are strongly affected by sequence. The compression on short length scales followed by expansion provides a plausible explanation for recent measurements in condensates which shows abnormally large exponents *ν >* 0.5 in *in-vitro* systems [15]. That low-complexity sequence by themselves appear to show bulk microphases behavior hints at highly non-uniform melts for actual proteins. This corroborates recent observations of micelles in various systems [16, 17], and presence of heterogeneity in condensates [18]. The consequences of heterogeneity remains to be studied; specifically, one could hypothesize that bulk microphase kinetically hinder fibrillation processes, e.g. by preventing contact between hydrophobic residues of different chains.

The non-linearity of patterning should also be used to constrain the protein predictors we use. Specifically, this article makes it clear that two parameters are needed, one related to mean-field, and one related to patterning. The current state-of-the-art predictor, the sequence hydropathy decorator [9], therefore does not have the appropriate functional form. This provides an explanation for its poor performance observed in [10]. The formalism deployed in this article is applicable to charge patterns, provided a regular screened electrostatics picture. A basic analysis (see SI) indicates that the mean-field picture is different and likely requires three parameters for a full characterization. These are expected to be salt-dependent, and their interplay to be responsible for the typical re-entrant phase separation of polyampholytes with salt [19]. Exact behavior of charge patterns, in particular for polymer melts, remains unclear at this point. The framework introduced here allows for fast estimation of sequence pattern effects. Using post-processing methods, the reweighting framework is of order 𝒪 (10^4^) faster than the direct simulation. On-the-fly estimation of partial derivatives of equation 2 could also provide an additional speed up. It can also be extended to tackle dynamical quantities, e.g. viscosity, through Girsanov reweighting [20]. However, it should be pointed out that the method requires high-quality data, in particular for long polymers, which may limit its applicability.

## METHODS

To determine single chain properties, we run simulations of 64 independent chains for each realization of the sequence, using the HOOMD-Blue molecular dynamics package [21–24].

We use a timestep Δ*t* = 0.01*τ*, where *τ* = *σ*ℳ ^1*/*2^*/ε*^1*/*2^ is the natural time unit of the system. Single chain simulations run for *t* = 5 · 10^5^*τ* . The base simulation length for melts is *t*_*M*_ = 2 · 10^6^*τ* . For *N* ≤ 128, simulations are at run for *t*_*M*_ . For *N* = 256, *ϕ* ≤ 0.3 and *N* = 512, *ϕ* ≤ 0.2, the simulations are run for 2*t*_*M*_ . Other cases are run for 4*t*_*M*_ .

All beads have an equal mass of ℳ. A Langevin thermostat with friction constant *γ* = 0.10 *ℳ/τ* is applied to obtain a canonical ensemble. The slopes of the cutting word are chosen randomly (uniform distribution). The sticker fraction is a by-product of the slopes, and is therefore also randomly distributed. The resulting distribution of *f* peaks near *f* = 1*/*2, and becomes sharper for larger ℵ.

Cutting words are created using a custom python package gitlab.mpcdf.mpg.de/mgirard/Words. Partial derivatives used to compute non-linear susceptibilities are computed using a custom package gitlab.mpcdf.mpg.de/mgirard/FastReweight. The versions used for the current article are also available as supplementary data. Molecular topologies are assembled using hoobas [25]. Covariances are computed using an online algorithm to minimize the memory footprint of the analysis. Analysis is performed using the freud package [26]. Computational workflow was managed by the signac-flow package [27–29].

## DATA AVAILABILITY

The complete workflow is available as part of the supplementary information. The dataset arising from analysis of trajectories is available at https://doi.org/10.17617/3.FZYS1O. Trajectories are available under reasonable request.

## ACKNOWLEDGEMENTS

I am grateful for numerous important discussions on polymer theory with Kurt Kremer and Burkhard Dünweg. Part of this work was motivated by numerous discussions on disordered proteins with Edward A. Lemke. I thank Jasper Michels and Kurt Kremer for a critical reading of this manuscript.

This work was supported by the Max-Planck Computing and Data Facilities, and financial support of the Collaborative Research Centre 1551 “Polymer concepts in cellular function” of the DFG (Deutsche Forschungsgemeinschaft, project number 464588647).

---

**Figure S1.**
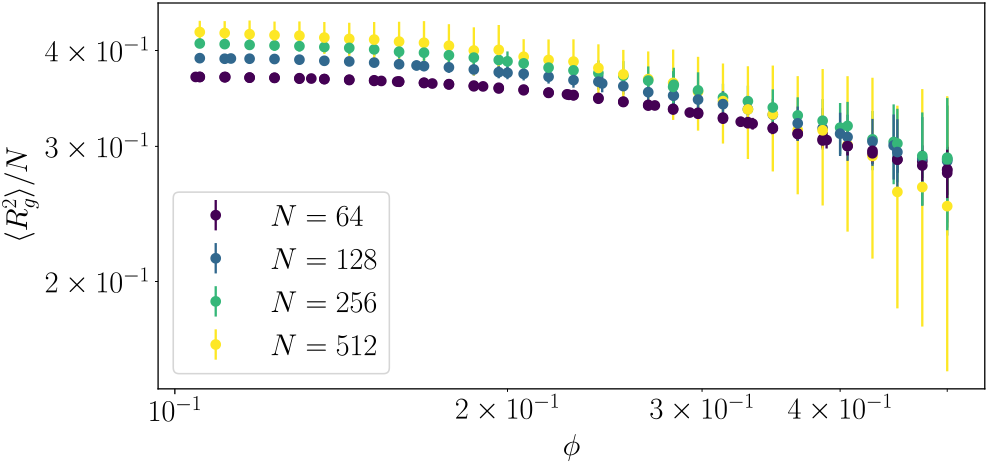
Radius of gyration of polymer chains in the melt. Unexpected behavior for *N* = 512 at large *ϕ* indicates equilibration issues. This is supported by the large error bars stemming from different chains exploring different portions of the conformational landscape.

## APPENDIX

### Equilibrating polymer melts

Equilibrating polymer melts is a notorious problem, complicated by the presence of glass transitions for polymer with attractive potentials. We refer the reader to various publications on the topic for further details []. The glass transition, and equilibration time, is tightly connected to the presence of entanglements. These are absent in our simulations by virtue of bond crossings. For long chains at high packing, this single measure is largely insu−icient to obtain fully equilibrated melts as shown on Fig. S1, where clear deviations are observed for *N* = 512 at large *ϕ*. The presence of large error bars indicate that the various chains have not sampled the same conformational landscape over the simulation time. For these particular samples, we find that increasing simulation time tends to increase the susceptibility observed.

Since susceptibility is computed from a mean-field, i.e. homopolymer, description, numerical issues associated with long chains may be alleviated by careful use of advanced Monte-Carlo moves involving bond swapping.

#### Sidechain decoration

We now consider an alternative disorder, namely in terms of the parameter *σ* along the chain, in analogous fashion to Δ. We now set *σ*_*A*_ = *σ*_0_ + *σ/f* and *σ*_*B*_ = *σ*_0_ − *σ/*(1− *f* ); and label this disorder *σ*. However, we have been unable to measure this response (see Fig. S3). Partial derivatives of the internal energy with respect to *σ* tend to yield large sparse outliers that are not well averaged over.

**Figure S2.**
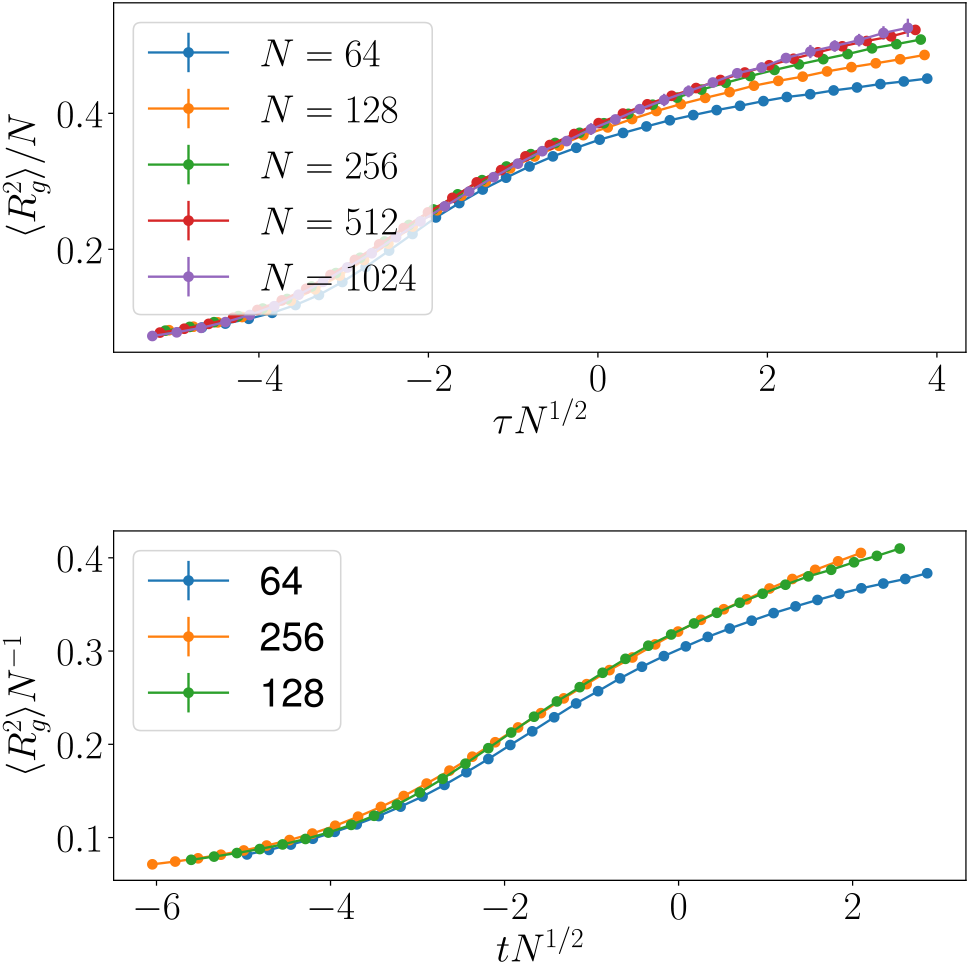
Finite-size scaling of the radius of gyration for a) homopolymers, showing collapse of the curves with *χ*_*θ*_ = 0.416 and b) heteropolymer (*ℵ* = 3, Δ = 0.2), showing some collapse of the curves for *χ*_*θ*_ *≈* 0.38. The two sets of curves show different values of 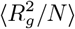 at *τ* = 0, namely *≈* 0.38 for homopolymers, and *≈* 0.30 for heteropolymers

**Figure S3.**
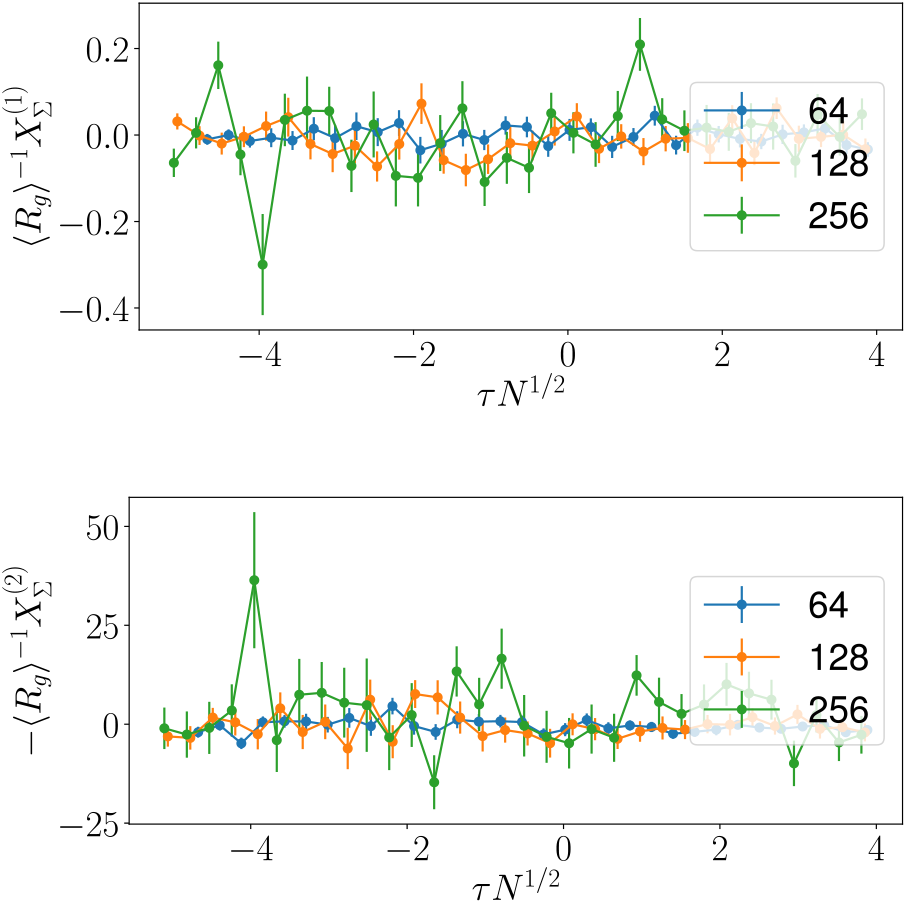
Susceptibility of the polymer chain versus the *σ* parameter of the Ashbaugh-Hatch potential. a) Linear response, and b) Quadratic response. Responses are below noise level of the measurements. Large deviations are caused by singular values tied to large partial derivatives of the potential with respect to *σ*.

**Figure S4.**
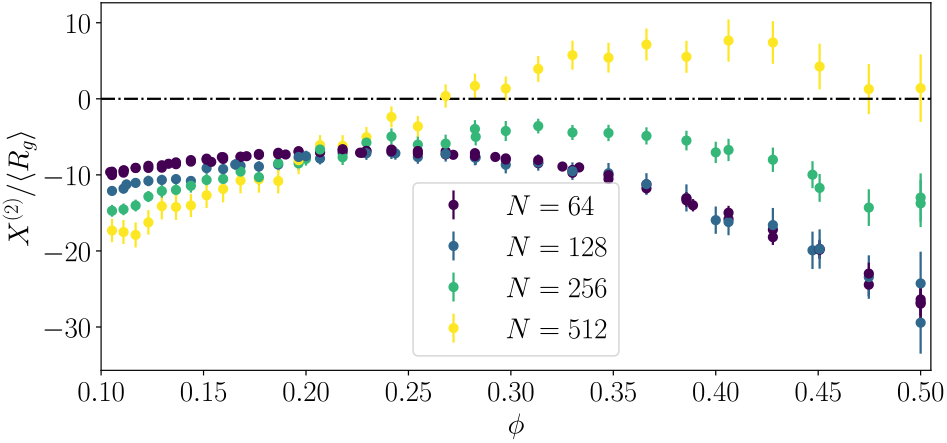
Susceptibility of polymer chains in a melt for *f* = 0.08

#### Charge patterns in the mean-field

Let us denote the fraction of charged monomer of each sign by *f* _±_, such that the total fraction of charged monomers is *f* = *f*_+_ + *f* _−_. For the convenience of the derivation, we assume here *f* = 1. Our starting point for a mean-field-less transformation is a polymer chain where each monomer carries charge *q*_0_ = *f*_+_ −*f*_−_ . The transformation for monomer is given by *q* = *q*_0_ ± *Q*(1 ∓ *q*_0_), where *Q* is the coordinate which imposes a charge patterning. Overall charge of the chain is fixed for the transformation, and the correct charge pattern is obtained for *Q* = 1. It is trivial to observe that the quantity *q*_0_ is essential to characterize the single chain behavior, in an analogous fashion to *χ*_*e*_ for our hydropathy decoration. Discarding the spatial part of charge interactions, the potential energy of a single contact would vary as:

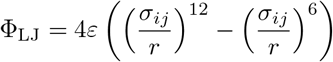

Given that the probability for a contact is the product of the fractions, we can easily compute ⟨*u*^(1)^ ⟩ = ⟨*u*^(2)^ ⟩ = 0, independently of *q*_0_. We can obtain the expected second order susceptibility with respect to *Q* as in our meanfield picture,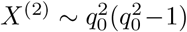. However, we now have an issue, since when *q*_0_ = 0 (charge neutral polymer),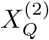 is expected to be zero. This needs to be reconciled with fact that sequence does play a role in charge neutral polymers [7]. Salvation comes from further expanding the series to *X*^(4)^, which includes terms in 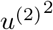.

This would suggest the relation between sequence and the susceptibilities of different orders are different. Therefore, three parameters would be needed to characterize a polyampholyte sequence: the net charge *q*_0_, *X*^(2)^ and *X*^(4)^. There is an analogy we can however make here, where the sequence introduces a “dipole”, and where *X*^(2)^ is a charge-dipole interaction, and *X*^(4)^ is a dipole-dipole interaction. This would imply that the two susceptibilities are related. Further investigation is clearly required, and in particular with respect to the effect of salt concentration.

## References

[1] S. F. Banani, H. O. Lee, A. A. Hyman, and M. K. Rosen, Biomolecular condensates: organizers of cellular biochemistry, Nature Reviews Molecular Cell Biology 18, 285 (2017), publisher: Nature Publishing Group.

[2] A. Bremer, M. Farag, W. M. Borcherds, I. Peran, E. W. Martin, R. V. Pappu, and T. Mittag, Deciphering how naturally occurring sequence features impact the phase behaviours of disordered prion-like domains, Nature Chemistry 14, 196 (2022), publisher: Nature Publishing Group.

[3] J. Wang, J.-M. Choi, A. S. Holehouse, H. O. Lee, X. Zhang, M. Jahnel, S. Maharana, R. Lemaitre, A. Pozniakovsky, D. Drechsel, I. Poser, R. V. Pappu, S. Alberti, and A. A. Hyman, A Molecular Grammar Governing the Driving Forces for Phase Separation of Prion-like RNA Binding Proteins, Cell 174, 688 (2018), publisher: Elsevier.

[4] N. B. Wilding, M. Müller, and K. Binder, Chain length dependence of the polymer–solvent critical point parameters, The Journal of Chemical Physics 105, 802 (1996).

[5] M. Rubinstein and R. H. Colby, Polymer physics (Oxford University Press, USA, 2003).

[6] G. L. Dignon, W. Zheng, R. B. Best, Y. C. Kim, and J. Mittal, Relation between single-molecule properties and phase behavior of intrinsically disordered proteins, Proceedings of the National Academy of Sciences 115, 9929 (2018), publisher: Proceedings of the National Academy of Sciences.

[7] R. K. Das and R. V. Pappu, Conformations of intrinsically disordered proteins are influenced by linear sequence distributions of oppositely charged residues, Proceedings of the National Academy of Sciences 110, 13392 (2013), publisher: Proceedings of the National Academy of Sciences.

[8] L. Sawle and K. Ghosh, A theoretical method to compute sequence dependent configurational properties in charged polymers and proteins, The Journal of Chemical Physics 143, 085101 (2015).

[9] W. Zheng, G. Dignon, M. Brown, Y. C. Kim, and J. Mittal, Hydropathy Patterning Complements Charge Patterning to Describe Conformational Preferences of Disordered Proteins, The journal of physical chemistry letters 11, 3408 (2020).

[10] M. Girard, Disordered proteins: microphases or associative polymers? (2024), pages: 2024.10.09.617362 Section: New Results.

[11] M. Lothaire, Combinatorics on Words (Addison-Wesley, Advanced Book Program, World Science Division, 1983).

[12] G. Tesei and K. Lindorff-Larsen, Improved predictions of phase behaviour of intrinsically disordered proteins by tuning the interaction range, Open Research Europe 2, 94 (2023).

[13] J. A. Joseph, A. Reinhardt, A. Aguirre, P. Y. Chew, K. O. Russell, J. R. Espinosa, A. Garaizar, and R. Collepardo-Guevara, Physics-driven coarse-grained model for biomolecular phase separation with near-quantitative accuracy, Nature Computational Science 1, 732 (2021).

[14] G. L. Dignon, W. Zheng, Y. C. Kim, R. B. Best, and J. Mittal, Sequence determinants of protein phase behavior from a coarse-grained model, PLOS Computational Biology 14, e1005941 (2018), publisher: Public Library of Science.

[15] M. Yu, M. Heidari, S. Mikhaleva, P. S. Tan, S. Mingu, H. Ruan, C. D. Reinkemeier, A. Obarska-Kosinska, M. Siggel, M. Beck, G. Hummer, and E. A. Lemke, Visualizing the disordered nuclear transport machinery in situ, Nature 617, 162 (2023), publisher: Nature Publishing Group.

[16] H. Ruan, R. Dillenburg, S. Wittmann, M. Girard, and E. A. Lemke, Differential conformational expansion of Nup98-HOXA9 oncoprotein in micro- and macrophases (2025), pages: 2025.05.21.655138 Section: New Results.

[17] M. K. Shinn, D. T. Tomares, V. Liu, A. Pant, Y. Qiu, Y. J. Song, Y. Ayala, G. W. Strout, K. M. Ruff, M. D. Lew, K. V. Prasanth, and R. V. Pappu, Nuclear speckle proteins form intrinsic and MALAT1-dependent microphases (2025), pages: 2025.02.26.640430 Section: New Results.

[18] T. Wu, M. R. King, Y. Qiu, M. Farag, R. V. Pappu, and M. D. Lew, Single-fluorogen imaging reveals distinct environmental and structural features of biomolecular condensates, Nature Physics 21, 778 (2025), publisher: Nature Publishing Group.

[19] P. G. Higgs and J. Joanny, Theory of polyampholyte solutions, The Journal of Chemical Physics 94, 1543 (1991).

[20] L. Donati, M. Weber, and B. G. Keller, A review of Girsanov reweighting and of square root approximation for building molecular Markov state models, Journal of Mathematical Physics 63, 123306 (2022).

[21] J. A. Anderson, C. D. Lorenz, and A. Travesset, General purpose molecular dynamics simulations fully implemented on graphics processing units, Journal of Computational Physics 227, 5342 (2008).

[22] J. A. Anderson, E. Jankowski, T. L. Grubb, M. Engel, and S. C. Glotzer, Massively parallel Monte Carlo for many-particle simulations on GPUs, Journal of Computational Physics 254, 27 (2013).

[23] M. P. Howard, A. Statt, F. Madutsa, T. M. Truskett, and A. Z. Panagiotopoulos, Quantized bounding volume hierarchies for neighbor search in molecular simulations on graphics processing units, Computational Materials Science 164, 139 (2019).

[24] J. A. Anderson, J. Glaser, and S. C. Glotzer, HOOMD-blue: A Python package for high-performance molecular dynamics and hard particle Monte Carlo simulations, Computational Materials Science 173, 109363 (2020).

[25] M. Girard, A. Ehlen, A. Shakya, T. Bereau, and M. O. de la Cruz, Hoobas: A highly object-oriented builder for molecular dynamics, Computational Materials Science 167, 25 (2019).

[26] V. Ramasubramani, B. D. Dice, E. S. Harper, M. P. Spellings, J. A. Anderson, and S. C. Glotzer, freud: A software suite for high throughput analysis of particle simulation data, Computer Physics Communications 254, 107275 (2020).

[27] C. S. Adorf, P. M. Dodd, V. Ramasubramani, and S. C. Glotzer, Simple data and workflow management with the signac framework, Computational Materials Science 146, 220 (2018).

[28] signac: A Python framework for data and workflow management - SciPy Proceedings (2018).

[29] signac: Data Management and Workflows for Computational Researchers - SciPy Proceedings (2021).

